# RefSeq database growth influences the accuracy of *k*-mer-based species identification

**DOI:** 10.1101/304972

**Authors:** Daniel J. Nasko, Sergey Koren, Adam M. Phillippy, Todd J. Treangen

**Affiliations:** Center for Bioinformatics and Computational Biology, University of Maryland, College Park, Maryland, USA.; Genome Informatics Section, Computational and Statistical Genomics Branch, National Human Genome Research Institute, Bethesda, Maryland, USA.

**Keywords:** Taxonomic classification, Reference database, Metagenomics, Microbiome, Comparative analysis

## Abstract

Accurate species-level taxonomic classification and profiling of complex microbial communities remains a challenge due to homologous regions shared among closely related species and a sparse representation of non-human associated microbes in the database. Although the database undoubtedly has a strong influence on the sensitivity of taxonomic classifiers and profilers, to date, no study has carefully explored this topic on historical RefSeq releases and explored its impact on accuracy. In this study, we examined the influence of the database, over time, on *k*-mer based sequence classification and profiling. We present three major findings: (*i*) database growth over time resulted in more classified reads, but fewer species-level classifications and more species-level misclassifications; (*ii*) Bayesian re-estimation of abundance helped to recover species-level classifications when the exact target strain was present; and (*iii*) Bayesian reestimation struggled when the database lacked the target strain, resulting in a notable decrease in accuracy. In summary, our findings suggest that the growth of RefSeq over time has strongly influenced the accuracy of *k*-mer based classification and profiling methods, resulting in different classification results depending on the particular database used. These results suggest a need for new algorithms specially adapted for large genome collections and better measures of classification uncertainty.

## INTRODUCTION

Fundamental questions of a metagenomic survey are: (*i*) what microbes are present in each sample, (*ii*) how abundant is each organism identified in a sample, (*iii*) what role might each microbe play (i.e. what gene functions are present) and (*iv*) how do the previous observations change across samples and time? Specifically, there have been numerous studies highlighting the utility of metagenomic datasets for pathogen detection, disease indicators, and health ^1,2^. Addressing each of these fundamental questions is predicated on the ability to assign taxonomy and gene function to unknown sequences.

Several new tools and approaches for taxonomic identification of DNA sequences have emerged ^3–5^, in addition to community-driven ‘bake-offs’ and benchmarks ^6^. *k*-mer based classification methods such as Kraken or CLARK ^3,7^ are notable for their exceptional speed and specificity, as both are capable of analyzing hundreds of millions of short reads (ca. 100 base pairs) in a CPU minute. These *k*-mer based algorithms use heuristics to identify unique, informative, *k*-length subsequences (*k*-mers) within a database to help improve both speed and accuracy. A challenge for *k*-mer based classification approaches is that closely related species and strains often contain many identical sequences within their genomes. This challenge is typically addressed by assigning the query sequence with the lowest common ancestor (LCA) of all species that share the sequence. A comprehensive benchmarking survey indicated that Kraken offered the best F_1_ score (a measure considering both precision and recall) among the *k*-mer based taxonomic classifiers evaluated at the species level ^8^. Bracken, a Bayesian method that refines Kraken results, is capable of estimating how much of each species is present among a set of ambiguous species classifications by probabilistically re-distributing reads in a taxonomic tree ^9^. We thus selected Kraken and Bracken as representative tools from the genre of *k*-mer based classification methods. The focus on this study was not to examine a specific software tool, but rather to decouple the performance of *k*-mer based methods from the underlying database.

Available *k*-mer based methods for taxonomic identification and microbiome profiling rely on existing reference databases. While several investigations have examined the influence of contamination in specific database releases, and identified idiosyncrasies specific to a release ^10,11^, no study has examined the specific influence of perhaps the most popular database from which to build classification databases, the repository of sequenced and assembled microbes (RefSeq), across all releases of the database. Additionally, metagenomic classification and profiling tools are commonly compared to each other using simulated datasets on a fixed database, with leave-one-out analysis, but never compared to each other across recent trajectories in database growth. The aim of this study was to elucidate the influence of RefSeq database growth over time on the performance of *k*-mer based taxonomic identification tools.

## RESULTS

### RefSeq database growth

Since its release in June 2003 bacterial RefSeq, on average, has doubled in size (giga base pairs, Gbp) every 1.5 years (Fig. 1A), with the number of unique 31-mers in the database growing at a similar rate (Fig. 1B). A more recent release, bacterial RefSeq version 84 (released 9/11/2017), totaled over 700 Gbp of sequence data. The Simpson’s index of diversity is a metric with values between zero and one that reports the probability that two individuals randomly selected from a sample will not belong to the same species. Samples with a high Simpson’s index of diversity (i.e. closer to one) may be considered more diverse than those with low values (i.e. closer to zero). The diversity for each version of the bacterial RefSeq database increased until April 2013 where the Simpson’s index of diversity for each subsequent bacterial RefSeq release has trended downward (Fig. 1C). A slower growth is also seen in the number of new bacterial species in each RefSeq version, indicating many of the same species are being sequenced repeatedly (Fig. 1D).

**Figure 1:**
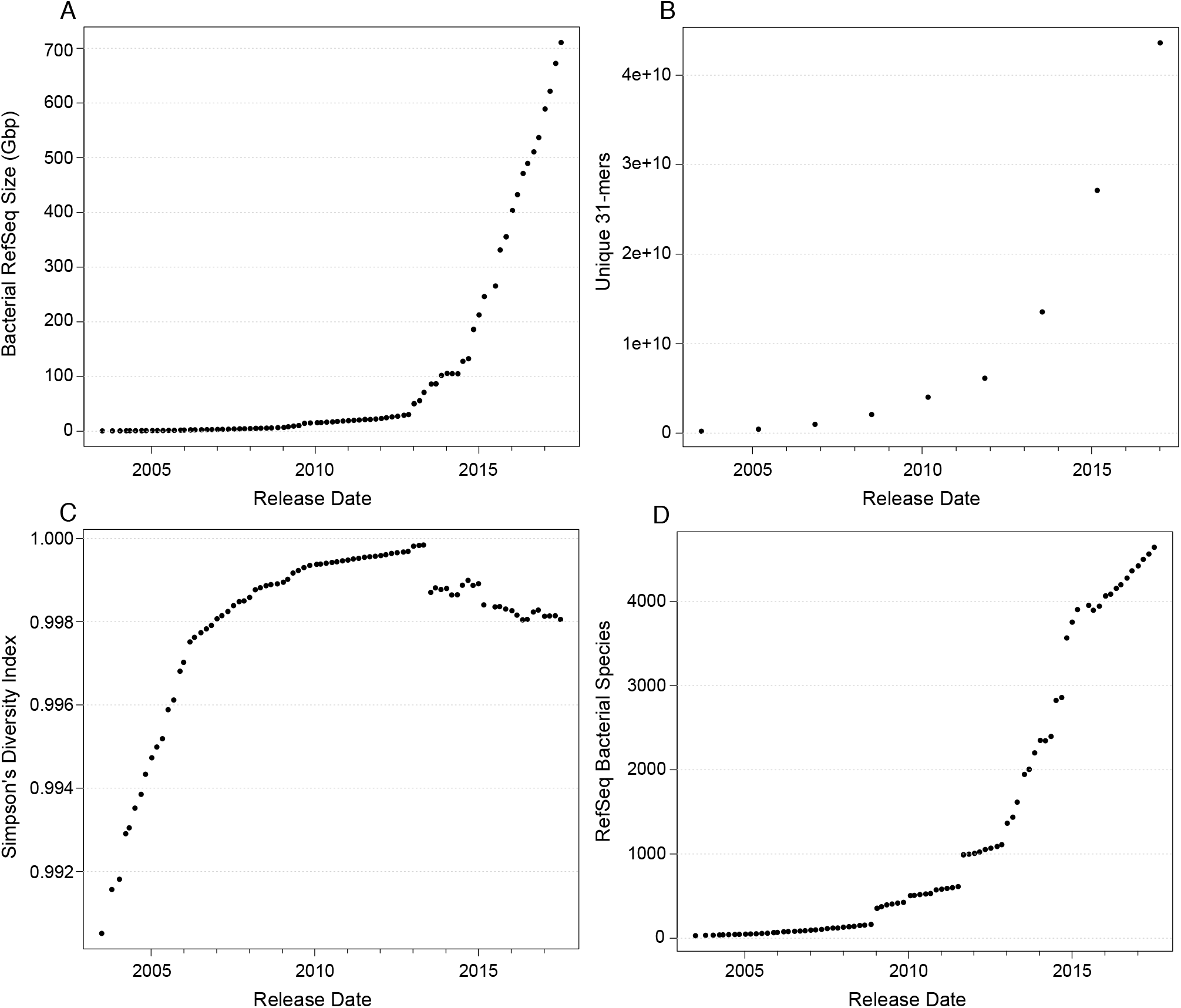
Simpson diversity index of bacterial RefSeq has decreased every release since April 2013. (A) The number of base pairs in bacterial RefSeq continues to grow exponentially, but (B) the number of unique 31-mers and (D) the number of bacterial species added increases slower. (C) The Simpson’s diversity index grew every release up to April 2013 where it has declined every release since.

### Taxonomic classification over time with a simulated metagenome

Kraken’s own simulated validation set of ten known genomes was searched against nine versions of bacterial RefSeq (1, 10, 20, 30, 40, 50, 60, 70, 80) and the MiniKraken database (4GB version) (Fig. 2). The accuracy of each Kraken run depends on the RefSeq version used in the search (Fig. 2; Table 1). Correct genus-level classifications increased as RefSeq grew, but correct species-level classifications peaked at version 30 and tended to decline thereafter (Fig. 2). The decrease in correct species classifications is due to more closely-related genomes appearing over time in RefSeq, making it difficult for the classifier to distinguish them and forcing a move up to the genus level. Overall, misclassified species-level calls were consistently rare, as reads were misclassified at the species level an average of 7% of the time (Table 1; Fig. 2). The fraction of reads classified at any taxonomic level, regardless of accuracy, increases as RefSeq grows over time (Fig. 3). However, the fraction of species-level assignments (again, regardless of accuracy) peaks at RefSeq version 30 and begins to decline thereafter, while the fraction of genus-level classifications begins to increase.

**Figure 2:**
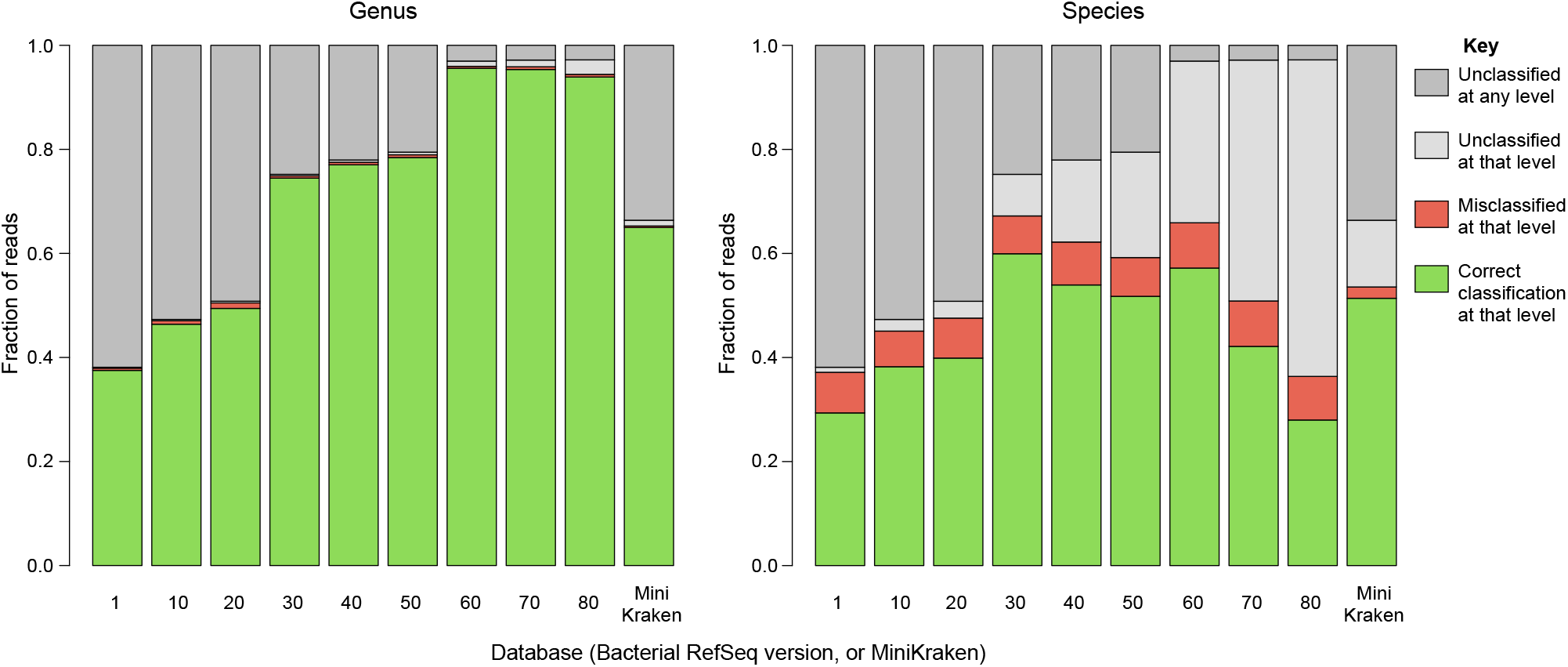
The fraction of correct species classifications (right) decreases in later RefSeq database versions because they are only being classified at the genus level (left). Kraken classification results of simulated reads from known genomes against nine versions the bacterial RefSeq database and the MiniKraken database. Misclassifications at the genus and species levels remain consistently low across database versions.

**Table 1.**
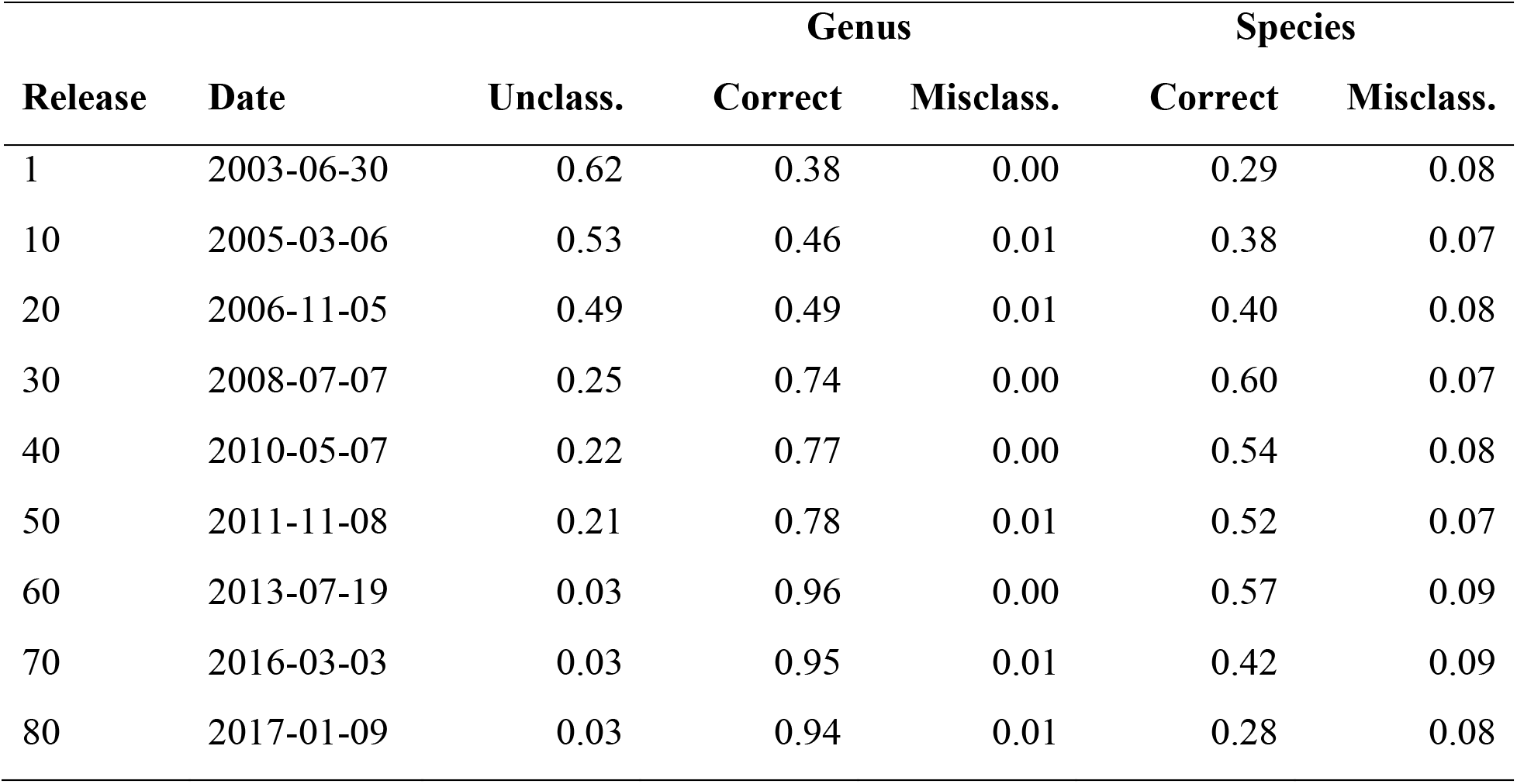
Fractions of unclassified (Unclass.), correctly classified (Correct), and misclassified (Misclass.) simulated reads from ten genomes using Kraken against different versions of bacterial RefSeq.

**Figure 3:**
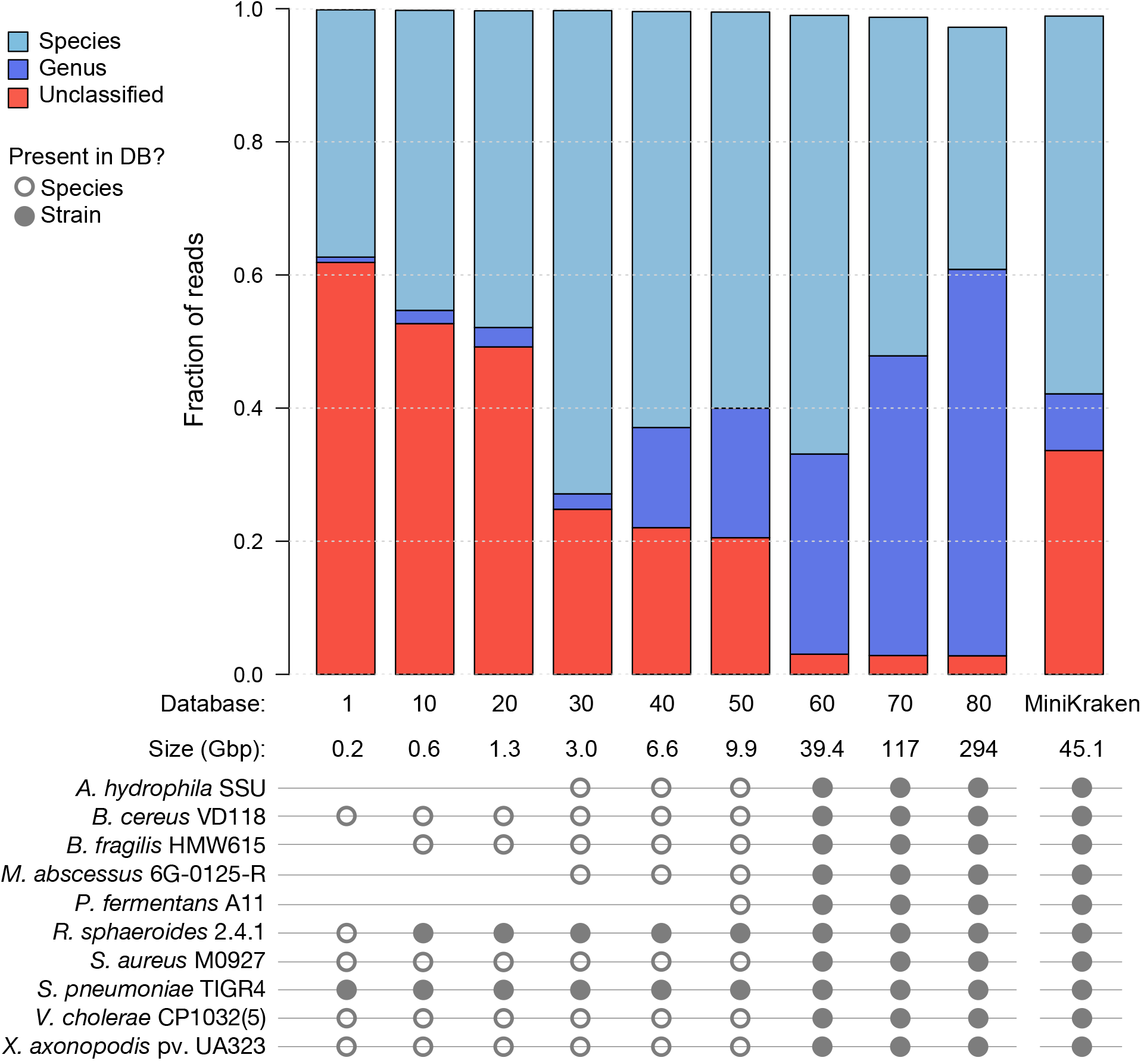
Species-level classifications decrease, and genus-level classifications increase as bacterial RefSeq grows. Fraction of simulated reads classified at different taxonomic levels, regardless of accuracy, using Kraken against ten databases. The circles below indicate when each genome’s species/strain is in a database. Although the MiniKraken database contains all 10 genomes it yields results comparable to bacterial RefSeq version 40.

Bracken was used to re-estimate the abundances of classifications made by Kraken when searching the simulated reads against eight bacterial RefSeq database versions (1, 10, 20, 30, 40, 50, 60, 70). Bracken first derives probabilities that describe how much sequence from each genome is identical to other genomes in the database. This step requires searching a Kraken database against itself with Kraken, which could not be performed for the MiniKraken DB (as there is no FASTA file for this database) or bacterial RefSeq version 80 (as it would require extensive computation for a database that size). Bracken was able to re-estimate species abundances for 95% of the input data using RefSeq version 70, while Kraken only classified 51% of reads at the species level. Because Bracken may probabilistically distribute a single read’s classification across multiple taxonomy nodes, its performance must be measured in terms of the predicted abundances. Bracken typically included the correct species in its re-estimation, but sometimes included incorrect species in the abundance estimation (on average 15% of reads were associated with a genome outside of the ten knowns).

### Taxonomic classification of difficult to classify genomes over time

The challenging nature of classifying sequences belonging to the *Bacillus cereus* sensu latu group has been previously documented ^12,13^. The *B. anthracis* species within this group is a well-defined monophyletic subclade of the larger *B. cereus* group, and the base of the *B. anthracis* clade is commonly denoted by a single nonsense mutation in the*plcR* gene ^14^ which is conserved in all known *B. anthracis* genomes and has been shown to confer a regulatory mutation essential for maintaining the pXO1 and pXO2 plasmids that carry the virulence factors characteristic of anthrax ^15^. However, not all *B. anthracis* cause disease in humans, such as *B. anthracis* Sterne (missing the pXO2 plasmid) and some *B. cereus* strains do cause anthrax-like disease ^16^, complicating a precise species definition. Thus, it is not a surprise that accurate species-level classification within this group has proven challenging for *k*-mer based methods, especially those methods not based on phylogenetic evidence. To demonstrate how difficult sequences from this group have been to classify over time, simulated reads were created for two *Bacillus cereus* strains. The first, *B. cereus* VD118, is a strain available in RefSeq version 60 and beyond, and the second, *B. cereus* ISSFR-23F ^17^, was recently isolated from the International Space Station and is not present in any of the RefSeq releases tested. It is phylogenetically close to *B. anthracis*, but lacks the phylogenetic and species characteristics of *B. anthracis.* Again, as bacterial RefSeq grows over time, the number of genus-level classifications made by Kraken increases (Fig. 4). While the number of genus-level calls made by Kraken increases over time the number of unclassified and misclassified species calls decreases (most commonly *B. anthracis, B. thuringensis*, and *B. weihenstephanensis).*

**Figure 4:**
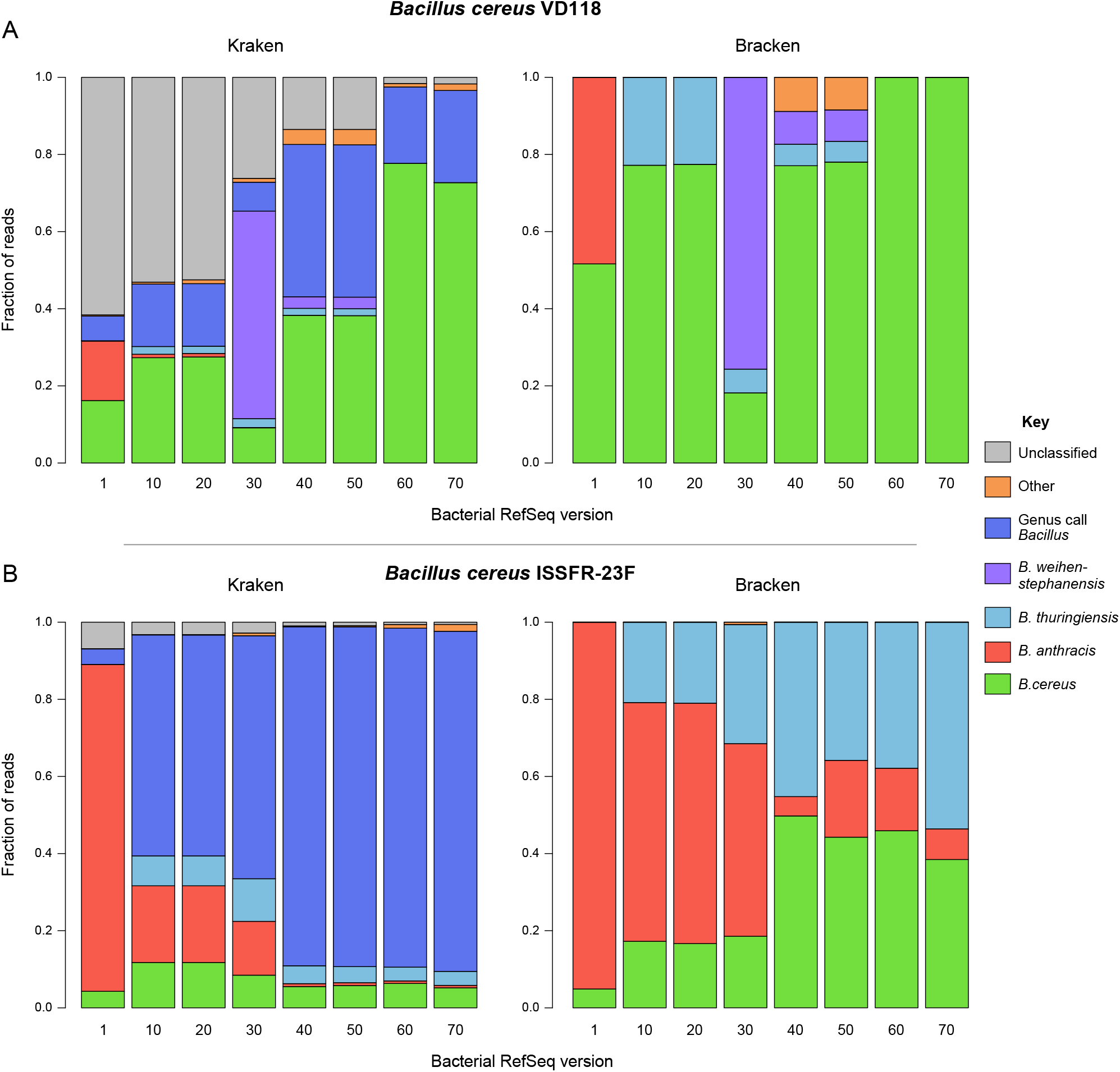
The fraction of simulated reads classified among *Bacillus* species varied considerably depending on which RefSeq version was used, demonstrating the influence of the database on a *k*-mer based taxonomic classification. (A) Classifying simulated *B. cereus* VD118 reads with Kraken (left) and Bracken (right) against different version of RefSeq. Species-level classifications varied, and the fraction of unclassified reads decreased with Kraken, as the database grew. Once *B. cereus* VD118 appeared in the database (ver. 60) Bracken correctly classified every read. (B) Species-level classifications decrease with Kraken as RefSeq grows using simulated reads from an environmental *Bacillus cereus* not in RefSeq. Fraction of simulated *B. cereus* ISSFR-23F reads classified using Kraken ver. 1.0 (left) and Bracken ver. 1.0.0 (right) against different versions of bacterial RefSeq. Bracken classification pushed all reads to a species-level call, though these classifications were often for other *Bacillus* species.

Bracken made species-level predictions for all reads no matter which version of bacterial RefSeq was used (Fig. 4). However, the increased rate of species-level predictions came at the cost of accuracy, as Bracken correctly identified *B. cereus* VD118 and *B. cereus* ISSFR-23F an average of 72% and 29% of the time, respectively, across RefSeq versions 1 through 70. The fraction of reads assigned to each *Bacillus* species varied substantially from each database tested. The range of *Bacillus* species predictions for *B. cereus* VD118 were: *B. cereus* 81% (max=100%, min=18%), *B. anthracis* 48% (max=48%, min=0%), *B. thuringiensis* 23% (max=23%, min=0%), and *B. weihenstephanensis* 76% (max=76%, min=0%). While the range of *Bacillus* species predictions for *B. cereus* ISSFR-23F were: *B. cereus* 45% (max=50%, min=5%), *B. anthracis* 90% (max=95%, min=5%), and *B. thuringiensis* 54% (max=54%, min=0%).

### CPU/Memory performance over time

Historical bacterial RefSeq versions were recreated and used to build Kraken databases with default settings. While most databases were constructed with ease and in less than a day, version 70 required 500 GB of RAM and 2 days (single compute node using on 64 cores), while version 80 required ca. 2.5 TB of RAM and ca. 11 days (single compute node using on 64 cores). Given this trend, future releases will likely require over 4 TB of RAM and weeks of computation to build, putting into question the feasibility of building and profiling *k*-mer databases on future RefSeq versions. Recent studies ^18^ have suggested alternative approaches for database construction that would help to circumvent future computational bottlenecks.

## DISCUSSION

The results of our study support three conclusions: (*i*) the RefSeq bacterial database composition and diversity is dynamic, varying from release to release; (*ii*) the database composition strongly influences the performance of *k*-mer based taxonomic identification methods, and (*iii*) Bayesian based methods can help mitigate some of the effect, but struggle with novel genomes that have close relatives in the database.

### Database influences on *k*-mer based taxonomic classification

Using Bracken, the majority of *Bacillus cereus* ISSFR-23F simulated reads were not correctly assigned to *B. cereus* but were more frequently mis-assigned as *Bacillus anthracis* or *Bacillus thuringiensis* (Fig. 4B). This, in part, is not surprising as two of the three species in this group, *B. cereus* and *B. thuringiensis*, have no clear phylogenetically defined boundary, though *B. anthracis* is phylogenetically distinct from *B. cereus* and *B. thuringiensis*. Furthermore, any two genomes within the *Bacillus cereus* sensu lato group are likely to be over 98% identical ^9^. Given that *k*-mer based methods are not phylogenetically-grounded, but rather based on sequence composition, they are susceptible to misidentification in clades where the taxonomy is in partial conflict with phylogeny, such as the *Bacillus cereus* sensu lato group. One clear example of misidentification within this group was the false identification Anthrax in public transit systems^19,20^

Another observation worth highlighting is that the fraction of simulated reads classified as one of the three *B. cereus* sensu lato species varied across database versions (Fig. 4), with the exception of *B. cereus* VD118, which was present in RefSeq releases 60 and 70 (Fig. 4A). The variation in species classifications across database versions indicates that even when using the same tools to analyze the same dataset, the conclusions derived from this analysis can vary substantially depending on which version of a database you are searching against, especially for genomes belonging to difficult to classify species (i.e. require phylogenetic-based approaches).

### Imperfect data

The genomic data deluge has helped to expand public repositories with a broader and deeper view of the tree of life, but has also brought with it contamination and misclassification. Contamination in public databases is well-documented ^21^ and represents an additional confounding factor for *k*-mer based methods. While several custom tools have been built to deal with imperfect data ^22^, there is a need for database ‘cleaning’ tools that can preprocess a database and evaluate it for both contamination (genome assemblies that contain a mixture of species) and misclassified species and strains (genomes that are assigned a taxonomic ID that is inconsistent with its similarity to other genomes in the database). The misclassification issue often is in the eye of the beholder; species have been named based on morphology, ecological niche, toxin presence/absence, isolation location, 16S phylogenetic placement, and average nucleotide identity across the genome. This, coupled with an often ambiguous species concept in microbial genomes due to horizontal gene transfer and mobile elements ^23^, brings into question the reliance on the current taxonomic structure for assigning names to microbes sequenced in metagenomic samples. A more robust approach would be for the classification databases to derive their own hierarchical structure directly from the data, rather than taxonomy, and then map back the internally derived hierarchy to widely-used taxonomic names.

## CONCLUSION

Our findings demonstrate that changes in RefSeq over time have influenced the accuracy of *k-* mer based classification and profiling methods. Bayesian re-estimation approaches are helpful for species or strain level prediction but can result in false positives and are computationally prohibitive with larger databases. Despite recent progress in *k*-mer based methods for metagenome profiling and classification these tools should likely be used as step one in a multi-step process, which also includes read mapping, assembly, feature prediction, and annotation. Additionally, priority should be given to the breadth, not depth, of species added to reference databases over time.

## METHODS

### Acquisition of bacterial RefSeq databases versions 1 through 80

FASTA files of previous versions of bacterial RefSeq are not publically available for download. Therefore, sequences from previous versions of bacterial RefSeq were acquired using custom scripts (https://github.com/dnasko/refseq_rollback). Briefly, the process involved downloading the current bacterial RefSeq release (ver. 84 as of the date of the analysis) FASTA files (ftp.ncbi.nlm.nih.gov/refseq/release/bacteria) and concatenating them into one file. Then, the catalog file associated with the desired version is downloaded(ftp.ncbi.nlm.nih.gov/refseq/release/release-catalog/archive), which contains the identifiers for sequences present in that version of bacterial RefSeq. Sequence identifiers in that version’s catalog file are pulled from the current RefSeq FASTA file and written to a new file. Using the refseq_rollback.pl script any version of bacterial RefSeq can be created. For this study only versions 1, 10, 20, 30, 40, 50, 60, 70, and 80 were recreated.

### Taxonomic classification on simulated datasets

Two simulated read datasets were used to test Kraken and Bracken performance with different versions of the bacterial RefSeq database. The first simulated dataset was downloaded from the Kraken website (ccb.jhu.edu/software/kraken), and was previously used in the Kraken manuscript as a validation set ^3^. Briefly, this simulated dataset was composed of 10 known bacterial species:*Aeromonas hydrophila* SSU, *Bacillus cereus* VD118, *Bacteroides fragilis* HMW 615, *Mycobacterium abscessus* 6G-0125-R, *Pelosinus fermentans* A11, *Rhodobacter sphaeroides* 2.4.1, *Staphylococcus aureus* M0927, *Streptococcus pneumoniae* TIGR4, *Vibrio cholerae* CP1032(5), and *Xanthomonas axonopodis* pv. Manihotis UA323. Each genome had 1,000 single-end reads (101 bp in size) for a total of 10,000 reads. We selected this dataset as it has been widely used as a benchmark for other *k*-mer based classification methods ^3,7^ and represents a breadth of species. This simulated read dataset was classified against each of the recreated bacterial RefSeq databases using Kraken (ver 1.0) with default settings.

To test the ability to classify reads from genomes not in the bacterial RefSeq database 10,000 simulated single-end Illumina reads (101 bp) were created using Grinder ^24^ with default settings from: (i) a *Bacillus cereus* genome, *B. cereus* VD118, not present in RefSeq until verion 60 and beyond; and (ii) a novel *B. cereus* genome, *B. cereus* ISSFR-23F ^17^, never present in any of the RefSeq versions tested. We decided to use these genomes as they are members of the *B. cereus* sensu lato group, containing a collection of species that are known to be challenging for k-mer methods to distinguish between ^19,20^. These datasets were classified with Kraken (ver. 1.0) and Bracken (ver. 1.0.0) ^9^ both with default settings (Bracken “read-length” set to 101).

### Running Bracken on Kraken output

Bracken (ver. 1.0.0) was run on the output of each Kraken search (except for release 80 and KrakenMiniDB). Default parameters were used except for “read-length”, which was set to 101.

### Bacterial RefSeq diversity metric calculations

Diversity metrics were calculated for every version of bacterial RefSeq (1-84) by parsing the catalog files for each version. An operational taxonomic unit (OTU) table was constructed using the NCBI taxonomy identifiers as taxonomic units (see create_otu_table.pl in the refseq_rollback repository). The OTU table was imported to QIIME (ver. MacQIIME 1.9.1-20150604) ^25^. Diversity metrics (Simpson, Shannon, Richness) were calculated using the “alpha_diversity.py” script and plotted using the R base package.

## ABBREVIATIONS

OTU: Operational taxonomic unit; LCA: Lowest common ancestor

## DECLARATIONS

### Acknowledgements

This work utilized the computational resources of the NIH HPC Biowulf cluster (https://hpc.nih.gov). The authors would like to thank Mihai Pop for his feedback and discussion of this project in its early development.

### Availability of Data and Materials

Scripts used in this analysis are available on GitHub

(github.com/dnasko/refseq_rollback). Datasets and genomes used in this analysis are available online and referenced in the text.

### Authors’ contributions

T.J.T. and D.J.N. designed the experiments. D.J.N. wrote the analysis scripts. D.J.N.,

S.K, and T.J.T. performed the experiments. D.J.N., S.K., A.M.P. and T.J.T. wrote the paper.

### Ethics approval and consent to participate

NA

### Consent for publication

NA

### Competing interests

NA

Additional files

